# Utilizing natural competence to genetically manipulate *Lactobacillus iners*

**DOI:** 10.64898/2026.03.03.709335

**Authors:** Kevin Y. Cao, Daniella Serrador, Jhenielle R. Campbell, Rupert Kaul, William Wiley Navarre

## Abstract

The healthy human vaginal microbiota is typically dominated by one species of *Lactobacillus*: *L. iners*, *L. crispatus*, *L. jensenii*, or *L. gasseri*. *L. iners*, the most prevalent vaginal microbe globally, is the most fastidious of the vaginal lactobacilli, has the smallest genome, and produces less lactic acid (only the L-isoform). *L. iners* is also less protective against bacterial vaginosis, and uniquely encodes a cholesterol-dependent cytolysin, inerolysin, suggesting it may be a pathobiont. Despite its central role in the health of over one billion females, *L. iners* biology remains poorly understood in part due to a lack of genetic editing tools. Here, we present findings that *L. iners* is naturally competent and can be transformed easily by exogenous DNA. Natural competence was leveraged to disrupt the *iny* gene encoding inerolysin, and *comGA*, encoding the ATPase component of the competence pilus. Both gene disruptions were accomplished using PCR assembled DNA fragments comprising a drug resistance gene cassette (*tetM* or *ermB*) flanked by ∼2 kb regions of homology to the *L. iners* chromosome. We further demonstrate that *comGA* is essential for *L. iners* transformation. The ability to rapidly perform targeted deletions in *L. iners* with *in vitro* generated DNA templates provides a straightforward and much needed method to probe the genetics and physiology of these important vaginal bacteria.

**IMPORTANCE:** This study describes, to our knowledge, the first method for genetically manipulating *L. iners*, the most prevalent bacteria of the human vaginal microbiota. This work paves the way for the rapid development of genetic tools to explore *L. iners*’ physiology in the context of the vaginal microbiome and potentially alter its properties as a probiotic.

## INTRODUCTION

The healthy human vaginal microbiome is typically dominated by a population of *Lactobacillus*: either *Lactobacillus crispatus*, *Lactobacillus jensenii*, *Lactobacillus gasseri*, or *Lactobacillus iners* (1). These vaginal lactobacilli are thought to afford protection against pathogenic anaerobic bacteria by producing lactic acid to acidify the vaginal microenvironment, as well as through production of bacteriocins (1).

*L. iners* is the most prevalent vaginal bacterium globally, and most individuals with *L. iners*-dominant microbiomes are healthy and remain stably colonized (2). *L. iners* is very fastidious, has a small genome of ∼1.3 Mbp, and does not encode genes necessary for the *de novo* synthesis of amino acids, suggesting a heavily host-dependent lifestyle (3, 4). Some *L. iners* strains also encode a lanthipeptide which may exert antimicrobial activity (5). However, individuals with an *L. iners*-dominated microbiome are more likely to transition to a dysbiotic state (bacterial vaginosis, or BV) than those with a *L. crispatus*-dominated microbiome (6). Furthermore, *L. iners* abundance is not significantly impacted during episodes of BV, and appears to not be displaced by BV-associated microbes like *Gardnerella* (6). *L. iners* is also the only vaginal *Lactobacillus* species to encode a cholesterol-dependent cytolysin, inerolysin, that can lyse epithelial cells (7). Virtually nothing is known about the regulation of inerolysin expression with one study finding it is upregulated in individuals with BV (8). This may suggest that *L. iners* is a weakly protective pathobiont that, under conditions of dysbiosis, upregulates a toxin that contributes to the pathology of BV.

To date, there have been no published reports of successful genetic editing of *L. iners*, and a previous attempt at gene editing *L. iners* using electroporation, conjugation and natural competence, was unsuccessful (9). Genetic tools for the other vaginal lactobacilli have been published including the use of an endogenous CRISPR system for editing *L. crispatus* (10), as well as heterologous CRISPR plasmids and specific base editing for *L. gasseri* (11, 12). *L. jensenii* (13) and *G. vaginalis* (14) have electroporation protocols to extrachromosomally express plasmid DNA. Electroporation of plasmids has also been used to gene-edit *G. vaginalis* (15).

Natural competence is found in both Gram-positive (e.g., *Bacillus subtilis*, *Streptococcus pneumoniae*, and *Streptococcus thermophilus*) (16) and Gram-negative bacteria (e.g., *Haemophilus influenzae*, *Neisseria gonorrhoeae* and *Neisseria meningitidis*) (17). In the lactic acid bacteria, the competence genes are controlled in a quorum sensing-dependent manner (16). A type IV-like pilus (encoded by the *comG* operon, with *comC* as a pilin subunit peptidase) (18–20) is required for initial binding and translocation of extracellular DNA to the cell membrane. Transport of DNA across the cellular membrane is mediated by a DNA uptake channel complex (encoded by *comE* and *comF* operons), in a process where one strand of DNA is degraded via an endA/nucA nuclease while the opposite ssDNA strand is translocated through the channel (16). Subsequent integration of the internalized ssDNA into the genome requires the general recombination functions encoded by *recA*, *dprA*, *ssbA/ssbB*, and the competence-associated factor, *coiA* (16).

Here we report that *L. iners* universally encodes the genes predicted to be sufficient for genetic competence and several strains can be transformed either with exogenously added chromosomal DNA purified from other strains of *L. iners*, or by short DNA cassettes assembled *in vitro* using PCR and Gibson assembly. *In vitro* generated DNA templates were successfully used to disrupt the inerolysin and *comGA* genes of *L. iners*, and the *comGA^−^* strains could no longer be transformed, confirming that the competence system is required for the genetic transformation we observe. We find drug-resistance genes encoded on *Tn*916 elements in some stains as useful markers for transformation and genetic manipulation, and leveraged Oxford Nanopore sequence read data to identify sites of DNA methylation across several strains. The ability to rapidly modify the chromosome of *L. iners* greatly expands our ability to explore its biology in the context of women’s health.

## MATERIALS AND METHODS

### Genomic analysis of competence in L. iners

Genome sequences of *L. iners* strains NVM020S14, NVM025S01, NVM041S27, NVM076S08, NVM196S05, and NVM210S01 were previously reported (21, 22) (BioProject Accession: PRJNA1184769). The GenBank reference genomes *L. iners* C0322A1 (RefSeq Accession: GCF_011058775.1), *Bacillus subtilis* 168 (GCF_000009045.1), *Streptococcus pneumoniae* TIGR4 (GCF_040687945.1), and *Streptococcus thermophilus* LMD-9 (GCF_000014485.1) were also used in analysis. Open reading frames corresponding to competence related genes in these strains are listed in Table S1.

BLASTP and TBLASTN searches were performed on competence genes in *B. subtilis*, *S. pneumoniae*, and *S. thermophilus* to compare to putative competence genes found in *L. iners*. Queries were done against both the non-redundant protein sequences (nr) and the nucleotide collection (nr/nt). Any hits were either BLAST against *L. iners* or were found from other vaginal *Lactobacillus* sp. and then BLAST against *L. iners*.

Annotated RefSeq sequences were downloaded based on BLAST results and clinker (23) was used to visualize clustering between each competence operon in the *L. iners* strains and the competent bacteria references.

For the phylogenomic analysis, across all strains, each competence gene was translated and underwent multiple sequence alignment using MAFFT (24) (parameters: LINSI model, --maxiterate 1000 –globalpair). Subsequently, alignments for each competence operon were concatenated per strain. The concatenated alignment was processed using IQ-TREE (25) to infer a maximum likelihood tree using a full partition/gene merge model (26, 27) and ultrafast bootstrapping (28) (-m MFP+MERGE -B 1000). The tree was visualized using iTOL (29).

### Transformation assay

*L. iners* strains used here are previously described (21). In brief, they were isolated from cervicovaginal secretions from Kenyan women with *L. iners*-dominated microbiomes. Media was deoxygenated for 24 hours before use in an anaerobic chamber (AS-580, Anaerobe Systems) using a gas mixture of 10% CO_2_, 10% H_2_, and balance N_2_ (Linde) (21). Glycerol stocks of *L. iners* strains were struck on New York City III agar supplemented with 10% heat-inactivated horse serum (Corning, 35-030-CV or Gibco, 26050088) plates (21, 30) and incubated anaerobically at 37 °C for 48 hours.

After incubation, 4-6 colonies from the agar plates were inoculated into 2 mL SLIM (21) and incubated at 37 °C in the anaerobic chamber for 24 hours. Overnight cultures were resuspended and subsequently diluted (in SLIM) to an optical density at 600 nm (OD_600_) of 0.5 using the Ultrospec 3100 Pro spectrophotometer (Amersham Biosciences). The diluted culture was then subcultured 1:100 in SLIM into a final volume of 200 μL in a 96-well plate (Sarstedt, 83.3924). The culture was incubated in the anaerobic chamber at 37 °C for 3 hours prior to adding DNA substrate. DNase controls had 0.5 µL of reconstituted DNase I (BioRad, 7326828 – powder was reconstituted in 250 µL 10 mM Tris, pH 7.5) added immediately prior to adding DNA substrate (detailed in Figure S3).

Genomic DNA isolation from *L. iners* strains was described previously; genomic DNA was extracted using the Promega Wizard Genomic DNA Purification Kit (Catalog #: A1120/A1125) (22); in addition, during the 600 µL isopropanol wash, 66.67 µL of 3 M, pH 5.2 sodium acetate was added to the gDNA mixture. Genomic DNA was quantified using the Invitrogen Qubit dsDNA BR Assay Kit (Q32850) and the Invitrogen Qubit 3 Fluorometer (Q33216). 100 ng gDNA was added to the subcultures. PCR knockout fragments were quantified using the NanoDrop ND-1000 Spectrophotometer, and 200 ng of PCR amplified DNA was added to the subcultures.

After addition of the DNA substrate (+/− DNase), the subcultures were incubated for 24 hours in the anaerobic chamber at 37 °C. To isolate transformants, 100-150 µL of the subcultures were spread onto NYC III agar plates with antibiotics. Plates contained 0.0625 µg/mL erythromycin or 0.625 µg/mL tetracycline. To quantify transformation efficiency, subcultures were serially diluted using sterile PBS, and 5-7.5 µL of the dilutions were spotted onto NYC III agar plates. Plates were incubated anaerobically at 37 °C for 48-72 hours and colony forming units (CFUs) were then quantified.

For significance calculations, multiple t tests were used (parameters used were: unpaired experimental design, non-Gaussian distribution, Mann-Whitney tests between conditions, no multiple comparison correction).

### Gibson assembly of inerolysin and *comGA* knockout PCR products

All primers are listed in Table S2. For assembling the knockout fragments, three Phusion polymerase (Thermo Scientific, F530S or New England Biolabs, M0530S) amplified fragments were assembled into a linear fragment for DNA uptake. The fragments had ∼20-40 bp homology between each other. Linear fragments were purified using the GeneJET PCR Purification Kit (Thermo Scientific, K0701). Cleaned fragments were added in a 1:1:1 mass ratio (100 ng of each fragment) to 15 µL of 1.33x Gibson Assembly Reaction Mix (31) to a final volume of 20 µL. Assemblies were incubated at 50 °C for 1 hour. Gibson assemblies were PCR amplified with Phusion polymerase to increase concentration of fragments prior to transformation, followed by purification using the GeneJET PCR Purification Kit.

### PCR identification of gDNA transformants and PCR knockout transformants

All primers are listed in Table S2. Colony PCR with Taq polymerase (Invitrogen, 10342020) was used to confirm that NVM196S05 transformants with NVM041S27 gDNA took up the *ermB* cassette, and that the inerolysin and *comGA* knockout fragments were successfully integrated into NVM196S05 and NVM076S08 respectively. Colonies were resuspended in 10 µL ddH_2_O prior to addition to PCR reaction.

### Methylation and motif analysis of *L. iners* strains

Genome extraction and sequencing was previously performed (22). After sequencing, dorado (using the sup model) (v1.1.1) (32) was used to basecall modified bases (parameters: duplex sup,6mA,4mC_5mC), demultiplex (command: demux) and trim adaptors (command: trim). Barcoded reads for each *L. iners* strain were mapped back using built-in minimap2 functionality in dorado (command: dorado aligner, default settings). Using modkit (v0.5.1) (33), aligned reads were used to create summary tables of methylation calls (command: modkit pileup, default settings) which was then used to summarize potential motifs each *L. iners* strain may be methylating (command: modkit motif search –min-sites 30). Motifs with ≥50 highly-likely/modified occurrences in the reads (high occurrence is number of basecalls for a motif that pass a fraction modified threshold of 60%) underwent evaluation (command: motif evaluate --known-motif) to finalize the list of methylated motifs present in each strain. If there were redundant motifs, the most simplified and shortest motif sequence was used. To identify potential restriction-modification systems and cross-examine modkit-obtained motifs, MicrobeMod (34) was used on the *L. iners* genomes (FASTA format) (command: annotate_rm, default settings).

## Results

### *L. iners* competence genes

We previously isolated and sequenced six *L. iners* strains from Kenyan women (22) and, during genome annotation, noted that all strains encode multiple operons and genes that would indicate they are naturally competent. Competence genes had previously been noted in the genomes of *L. iners* strains C0011D1 and AB1 (3, 16). In Gram-positive bacteria, the genes required for competence are distributed across several unlinked operons including *coiA*, *comC*, *comK*, and the polycistronic operons *comE*, *comF*, and *comG* (Figure 1). A further analysis of all 84 complete *L. iners* RefSeq genomes (available from NCBI in July of 2025) revealed that competence genes are universally present in all strains of *L. iners*, with one exception being a metagenomic assembly lacking the *comG* operon (GCF_022456315.1), which may not represent the sequence of a true isolate. Across *L. iners* strains, the various competence-associated gene loci are highly conserved in both genomic location and sequence. One region of significant sequence variation between strains lies at the 3’ end of the *comG* operon where the small subunits of the competence pilus (ComGF/ComGG) are encoded.

**Figure 1.**
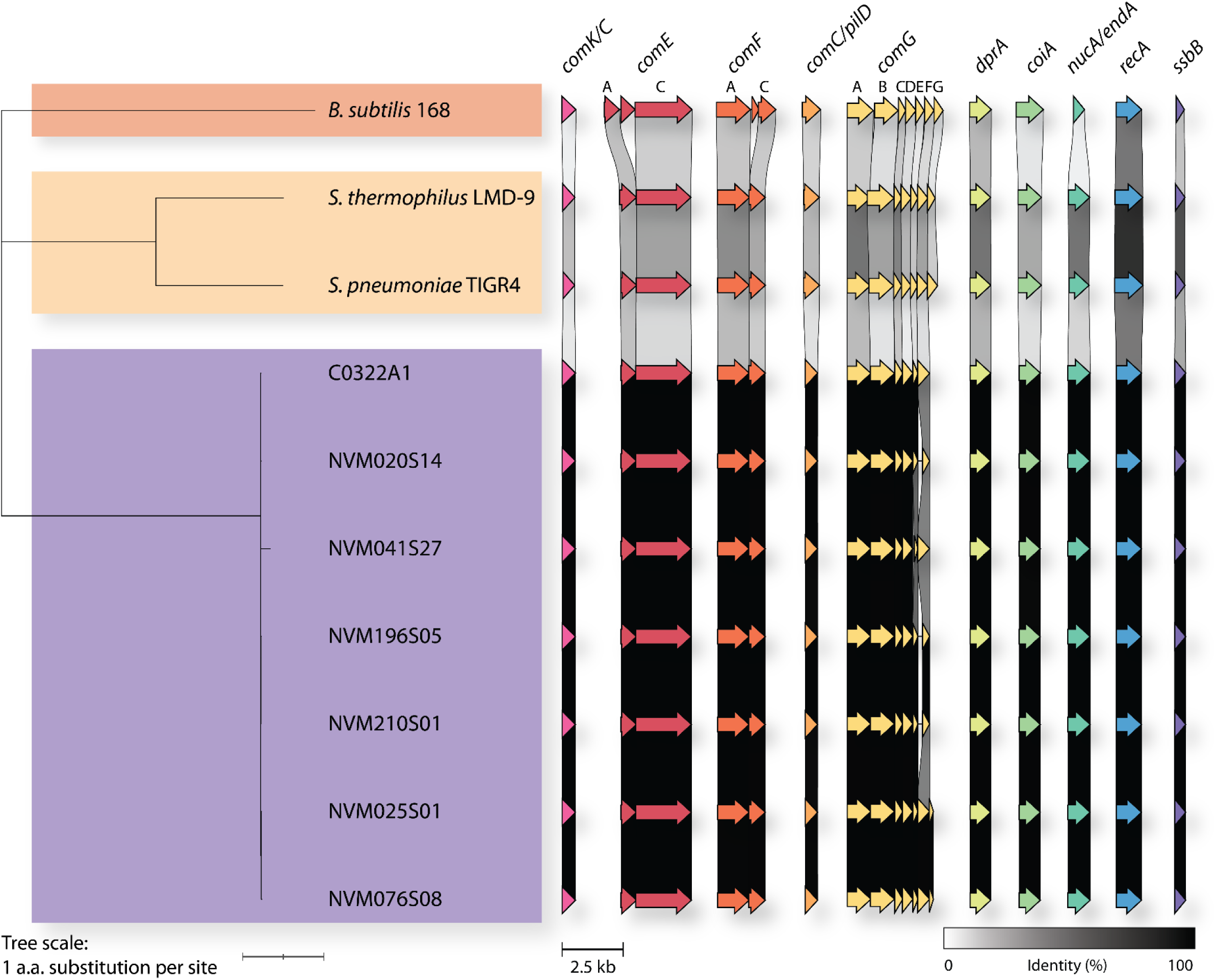
Genomic analysis of different *L. iners* strains to determine potential for natural competence. *L. iners* strains from Kenyan sex workers were examined for competence genes that may be homologous to model Gram-positive naturally competent bacteria (*B. subtilis*, *S. thermophilus*, *S. pneumoniae*). Competence genes across each strain/branch were concatenated and translated to protein for phylogenetic analysis. a.a. = amino acid. Clinker analysis (23) to right of tree shows nucleotide similarity between different competency genes.

A comparison of *L. iners* competence-associated genes against other naturally competent Gram-positive bacterial species such as *S. thermophilus*, *S. pneumoniae*, and *B. subtilis* showed the *comE*, *F* and *G* operon structures are largely syntenic between all these species (Figure 1, S1, Table S1). Protein sequence similarity between the competence-associated proteins of *L. iners*, *B. subtilis*, and the streptococci was often less than 40%, comparable to the similarity between functionally identical proteins across these genera.

### *L. iners* can be transformed with genomic DNA

Many *L. iners* isolates in our collection harbor a Tn*916* conjugative transposable island encoding tetracycline resistance via a *tetM* gene (Figure 2A and Figure S2). TetM, a GTPase structurally similar to the elongation factors EF-G and EF-Tu, binds the ribosome and hydrolyzes GTP to induce a conformational change that dislodges tetracycline from the 30S subunit (35). Tn*916* in two of our isolates also carry the *ermB* gene encoding ErmB (erythromycin ribosome methylase B) and its associated leader peptide ErmL. ErmB methylates the adenine residue (A2058) in the 23S rRNA of the 50S ribosomal subunit to confer resistance to erythromycin and other **m**acrolides, **l**incosamides, and **s**treptogramin **B** (the MLS-B phenotype) (36). Across the 84 sequenced RefSeq *L. iners* strains we analyzed, 52 isolates harbor Tn916 (all of which encode TetM), and 14 of those 52 strains have a Tn916 encoding both TetM and ErmB. When present, Tn916 is found in the same location in all *L. iners* strains, flanked by the *uspA* gene and a gene with homology to *ftsW/rodA*.

**Figure 2.**
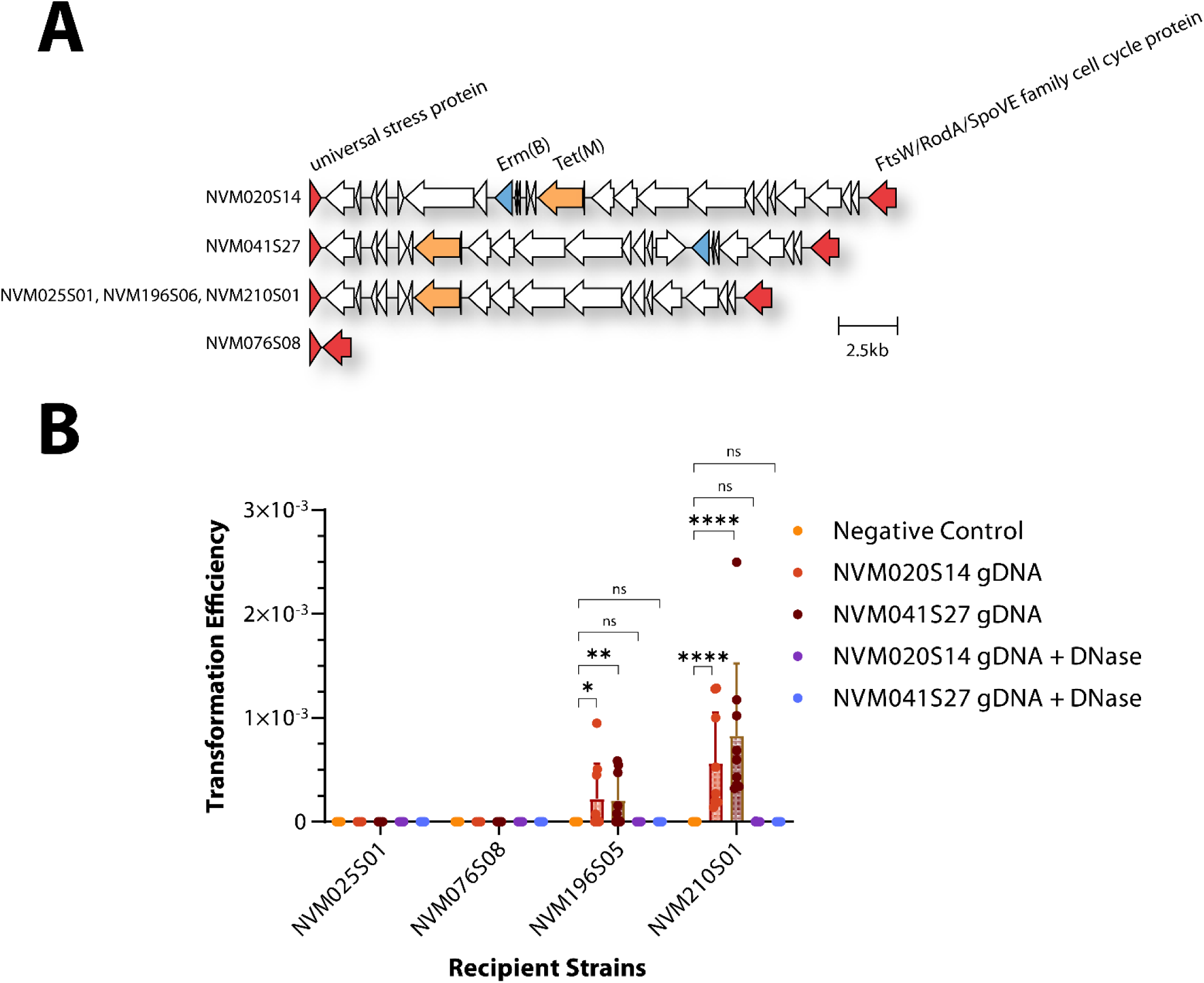
*L. iners* can be transformed using heterologous gDNA. **(A)** The distribution and arrangement of Tn*916* elements amongst *L. iners* strains in our collection, two of which carry both *tetM* and *ermB*. (**B)** Transformation efficiencies of erythromycin susceptible *L. iners* – calculated by taking ratio of colony forming units (CFU) of selected colonies (erythromycin-containing agar) divided by non-selected colonies (regular NYC agar) per respective condition. Mean transformation rate with NVM020S14 gDNA = 2.22 × 10^−4^ ± 3.39 × 10^−4^ for NVM196S05 and 5.64 × 10^−4^ ± 4.88 × 10^−4^ for NVM210S01; mean transformation rate with NVM041S27 gDNA = 2.06 × 10^−4^ ± 2.54 × 10^−4^ for NVM196S05 and 8.25 × 10^−4^ ± 6.98 × 10^−4^ for NVM210S01. ns = P > 0.05, * = P ≤ 0.05, ** = P ≤ 0.01, *** = P ≤ 0.001, **** = P ≤ 0.0001, N=3 (3 independent experiments), n=3

In susceptibility tests we find that strains carrying Tn916 elements are resistant to tetracycline at 0.625 µg/mL, and the subset of those strains that encode ErmB are also resistant to 0.0625 µg/mL erythromycin. One of our isolates, NVM076S08, does not harbor Tn*916* and, accordingly, we find it to be susceptible to both tetracycline and erythromycin (Figure 2A and Figure S2). This natural variation in drug resistance among our *L. iners* isolates provided us a straightforward way to determine if *L. iners* is naturally competent by testing if drug-susceptible strains acquire resistance when incubated with genomic DNA isolated from resistant strains.

Four erythromycin susceptible strains (NVM025S01, NVM076S08, NVM196S05 and NVM210S01) were grown in SLIM and then subcultured in a 1:100 dilution to grow for 3 hours prior to adding purified genomic DNA (gDNA, 100 ng) from two erythromycin-resistant strains of *L. iners* (NVM020S14 and NVM041S27, Figure S3). Cultures were incubated a further 24 hours before plating on New York City III (NYC III) agar with erythromycin. Resistant colonies were obtained from both NVM196S05 and NVM210S01, indicating they were successfully transformed (Figure 2B). Addition of 0.5 µL DNase to cultures immediately prior to the addition of gDNA significantly reduced or abrogated transformation (Figure 2B).

Transformation was confirmed by PCR. Primers were designed to anneal to recipient- or donor-specific genes to verify the genetic background of the isolate, and to *ermB* to verify that resistance was caused by acquisition of that gene. As shown in Figure 3, prior to transformation, the recipient strain NVM196S05 harbors an N^6^-methyltransferase, but lacks the *cas9* and *ermB* genes present in the donor strain NVM041S27. PCR of erythromycin-resistant colonies demonstrated that they retained the N^6^-methyltransferase gene, lacked the *cas9* gene (specific to the NVM196S05 genetic background), and had acquired the *ermB* gene. These results demonstrate that at least some strains of *L. iners* are naturally competent, that naked DNA is the transforming substrate and not phage or membrane enclosed DNA, and that the predicted erythromycin- and tetracycline-resistance genes encoded on the Tn*916* element found in many *L. iners* strains are active and confer resistance to the expected antibiotics.

**Figure 3.**
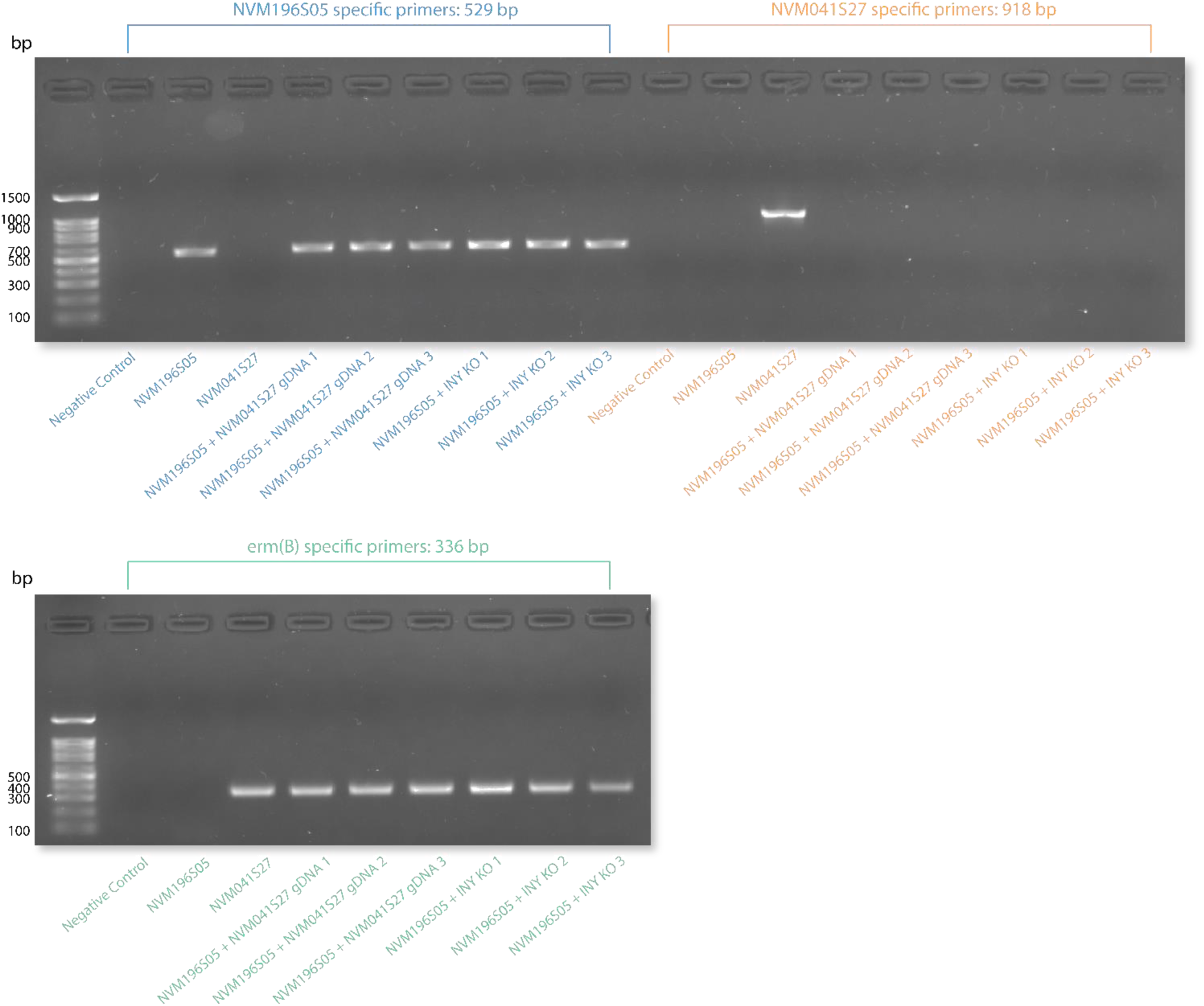
Colony PCR to differentiate between transformed and untransformed NVM196S05. n=3 (3 independent colonies) for experimental conditions. Negative control indicates no colony added. Expected primer sizes listed at top of gels.

### *L. iners* inerolysin knockout using PCR fragments

Given the demonstration of competence in two strains, an attempt was made to disrupt the *iny* gene encoding the pore-forming cytolysin, inerolysin (7), in our four erythromycin-susceptible strains. To do this, we PCR amplified two ∼2 kb DNA segments from the recipient strains, one including the 5’ ∼750 nucleotides of the inerolysin coding sequence (and ∼1250 bases upstream) and the other including the 3’ ∼750 nucleotides followed by another ∼1250 nucleotides downstream. We used the parental strain DNA as the PCR template for each transformation (i.e., the DNA from strain NVM196S05 was used to create the knockout cassette used for strain NVM196S05), because imperfect homology can significantly reduce the efficiency of recombination and transformation (37). We amplified the *ermB* gene including its putative promoter and leader peptide sequence from strain NVM041S27 and used Gibson assembly to generate synthetic DNA fragments where the erythromycin resistance segment is flanked on either side by ∼2 kb of DNA homologous to the *iny* gene.

We were able to transform 3 of the 4 recipient strains with these disruption cassettes (Figure 4). Again, colony PCR was used to confirm that for one of the recipient strains after knockout fragment addition, the transformants had the *ermB* cassette (Figure 3). In comparison to the gDNA transformations, NVM196S05 and NVM210S01 had lower transformation rates (Figure 4). However, NVM076S08 transformed successfully unlike what we observed above, when it was given gDNA from other strains and assessed for its ability to incorporate Tn*916*. It is possible that the region of DNA that needed to recombine was too large; NVM076S08 only has the *UspA* and *FtsW* flanking genes whereas the other recipients harbored a Tn*916* merely lacking *ermB*. It is also possible that sequence variation between NVM076S08 and donor strain DNA dramatically lowered recombination rates. By contrast, every strain has the inerolysin gene, and because the Gibson-generated knockout cassettes were amplified from the recipient strains’ DNA, transformation efficiency was improved. Finally, there may be differences in DNA modifications that limit strain-to-strain transformability. For example, NVM025S01 has a type I system that NVM076S08 lacks, which may be responsible for why transformants were not observed in that strain when using gDNA (Figure S4 and S5).

**Figure 4.**
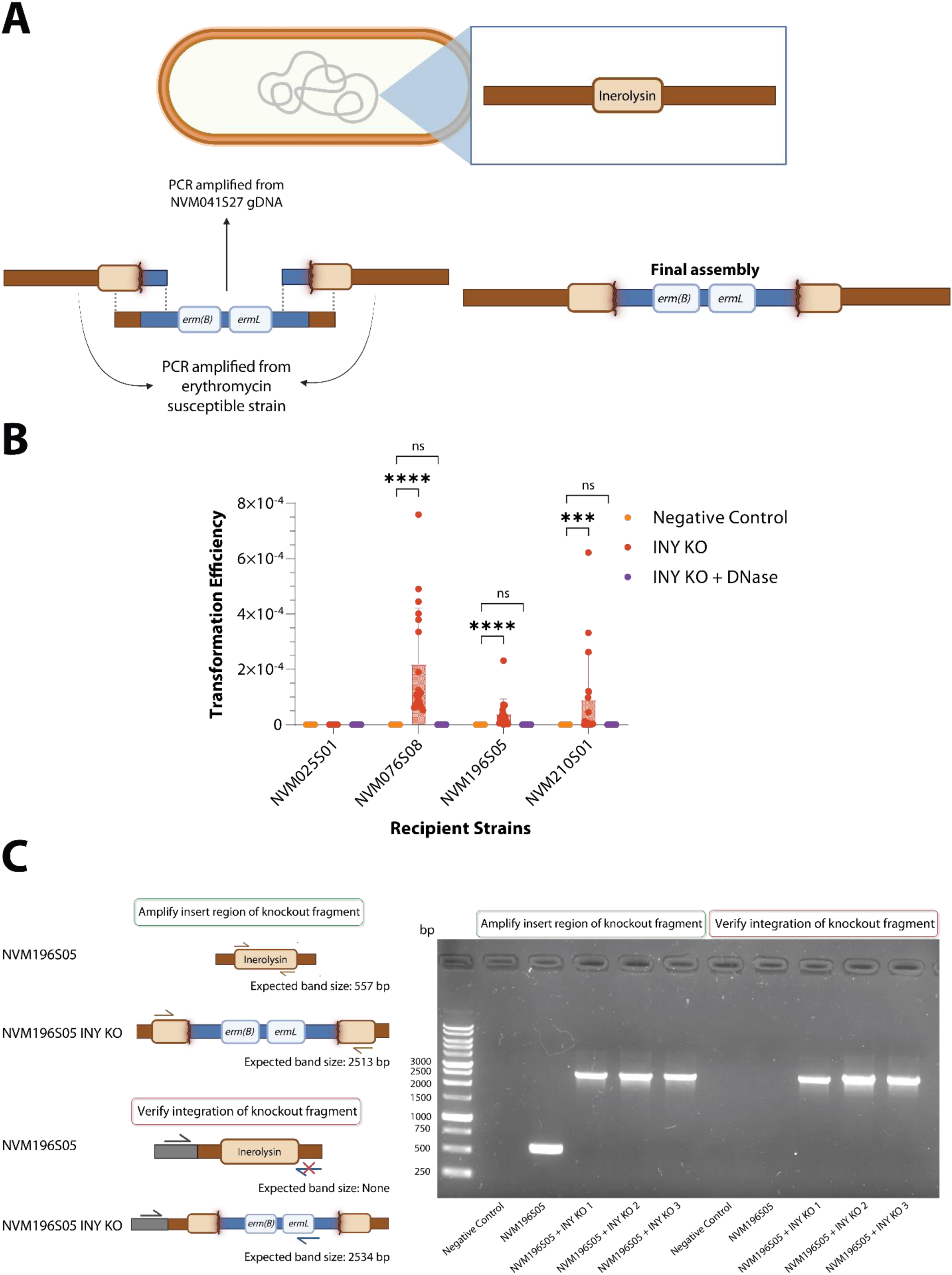
*L. iners* can be transformed with PCR fragments to create an inerolysin knockout. **A** – General overview of Gibson assembly to generate knockout fragments with upstream and downstream inerolysin fragments with homologies to each given erythromycin susceptible strain. **B** – Transformation efficiencies of erythromycin susceptible *L. iners* – calculated by taking ratio of colony forming units (CFU) of selected colonies (erythromycin-containing agar) divided by non-selected colonies (regular NYC agar) per respective condition. Mean transformation rate for NVM196S05 = 4.03 × 10^−5^ ± 5.33 × 10^−5^, 9.12 × 10^−5^ ± 1.64 × 10^−4^ for NVM210S01, and 2.19 × 10^−4^ ± 2.02 × 10^−4^ for NVM076S08. ns = P > 0.05, * = P ≤ 0.05, ** = P ≤ 0.01, *** = P ≤ 0.001, **** = P ≤ 0.0001, N=3 (3 independent experiments), n=3 for negative control, n=6 for INY KO and INY KO + DNase. **C** – Gel confirmation of knockout fragment in genome; insert region amplification verifies if there is an additional insert in the middle of the inerolysin gene and integration verification checks if fragment is inserted into genome by having one primer bind to middle of *ermB* cassette and one outside of fragment region (on genome itself). Negative control indicates no colony added.

### comGA is required for L. iners competence

To confirm that the competence system predicted bioinformatically is necessary for the transformation phenomenon we observe, and to demonstrate creation of a functional knockout, a disruption of one of the competence genes, *comGA*, was attempted. This gene encodes an ATPase responsible for assembly of the competence pilus, and is required for DNA uptake and transformation (20). *comGA* was chosen as it is most consistently annotated across *L. iners* genomes, and its disruption is predicted to eliminate transformation which gives us a functional assay to assess whether its knockout was successful.

The same Gibson assembly strategy was employed as for the inerolysin knockout, except using tetracycline resistance (*tetM*) from NVM041S27 as a selection marker. We chose NVM076S08 as our recipient strain as it had the highest PCR fragment transformation efficiency across our panel and it was the only fully sequenced strain in our isolate collection without native Tn*916*-encoded tetracycline resistance. Similar attempts to knockout *comGA* in NVM076S08 using a chloramphenicol resistance gene (*cmr*) amplified from plasmids pGT-Ah1 and pTRK669 have been unsuccessful. We do not know if this is because *cmr* is not functional in *L. iners* (to our knowledge, no wild *L. iners* have been observed to encode *cmr*), or if some sequence feature (hairpin structures, restriction motifs, etc.) reduces its ability to be acquired via competence.

As with the inerolysin disruption, we PCR amplified two fragments homologous to NVM076S08, one including the 5’ 501 nucleotides of *comGA* and 1,500 nucleotides upstream, and the other containing the 3’ 475 nucleotides of *comGA* and 1,500 nucleotides downstream (each ∼2 kb). We used Gibson assembly to combine these fragments to flank the *tetM* gene (with the 376 nucleotides upstream and 100 nucleotides downstream). After transformation, insertion was confirmed with colony PCR (Figure 5A). We then attempted to transform the *comGA* knockout with the inerolysin knockout fragment used previously. While NVM076S08 transformed as previously observed, the NVM076S08 *comGA^−^* strain did not produce erythromycin-resistant transformants (Figure 5B).

**Figure 5.**
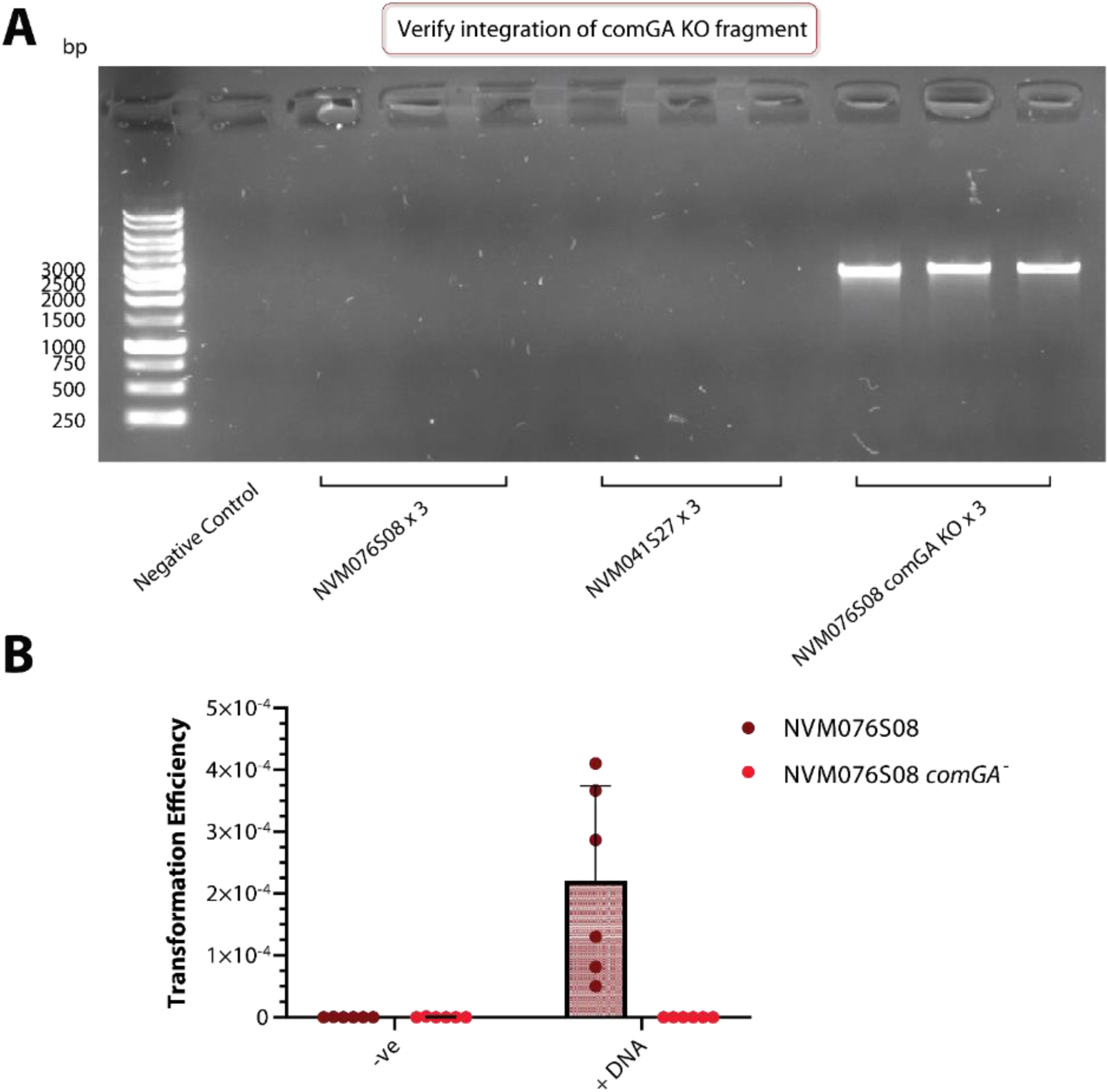
Knocking out *comGA* prevents *L. iners* transformation. **A** - Confirmation of *comGA* knockout using one primer targeted for *tetM* and one primer targeted to the genome upstream of the insert. Expected band size of 2663 bp. Negative control indicates no colony added. **B** - Transformation efficiency of *L. iners* NVM076S08 vs *comGA* knockout (NVM076S08 *comGA*^−^). Strains were transformed using the inerolysin knockout fragment with erythromycin resistance previously described. Ratio of CFUs on erythromycin NYC plates (selective) over CFUs on NYC plates (nonselective). N=2 (2 independent experiments), n=3.

To demonstrate that the lack of transformants was not due to NVM076S08 being unable to tolerate the disruptions in both genes or both antibiotic resistance markers, we reversed the order of events and transformed the NVM076S08 *iny::ermB* knockout strain using the *comGA::tetM* knockout fragment. This successfully generated the double knockout (Figure S6).

### DNA restriction-modification systems and methylations across *L. iners* isolates

The different transformation rates between strains like NVM210S01, which transformed easily, and NVM025S01, which did not yield transformants, could be caused by many factors including differences in regulation of the competence apparatus or by anti-DNA defense systems. An obvious possibility is that restriction-modification (RM) systems limit the rate of transformation across different strains of *L. iners*. MicrobeMod (34) was used to predict potential RM systems in each strain. All strains share an identical type II RM system predicted to modify GCGC motifs, as well as a conserved type I restriction enzyme (Figures S4 and S5). The two strains most readily transformed by gDNA, NVM196S05 and NVM210S01, lack a type III RM system and have few type I and type II restriction enzymes (Figure S4).

We previously sequenced the genomes of our isolates using Oxford Nanopore technology (22), which allowed us to retrospectively predict the presence or absence of specific DNA modifications through an analysis of electric current differences during sequencing (38). Nanopore data was analyzed with Modkit (33) to identify predicted methylation motifs from the raw sequence reads (Table S3). DNA modifications that can be inferred with this approach include 4- and 5-methylcytosine (4mC and 5mC respectively) and N6-methyladenine (6mA). Many, but not all, of the methylations observed by this method could be linked to the predicted specificity of a methylase encoded in the genome of the strain. All *L. iners* strains are extensively 5mC modified at the first cytosine within G[C]GC sequences, which can be explained by the universal presence of a type II RM system predicted to target GCGC motifs. Strain NVM041S27 displayed modifications at the sequence GGNN[C]C, which matches the predicted specificity of a type II methyltransferase in its genome. Strains NVM025S01 and NVM076S08 had extensive 6mA modifications at GC[A]GC sequences, in line with the predicted specificity of a type III methyltransferase they encode. NVM076S08 also encodes a type II restriction methylase with predicted CTKMAG specificity that corresponds to its abundant CTGC[A]G methylations. Many other modifications observed in the various isolates were not easily linked to a predicted methyltransferase. Our analysis reveals that *L. iners* strains harbor extensive methylations across their genomes that correspond largely, but imperfectly, with the predicted presence/absence of specific restriction-modification systems.

The role of RM systems with respect to natural transformation, if any, remains unclear. This initial report on the competence of *L. iners*, has only measured transformation at specific short regions of the genome (*iny*, *comGA* and *Tn*916) in a limited number of strains. We did not construct RM mutants, transform with DNA that has/lacks predicted target motifs, or with gDNA isolated from isogenic strains. We discuss potential sources of strain-to-strain variation in transformability further below.

## DISCUSSION

To our knowledge, we are reporting the first demonstration of natural competence in *L. iners* and its subsequent use for targeted genetic modification. Overall, the competence genes in *L. iners* strains are universally distributed and are highly similar across strains although some variability appears at the 3’ end of the *comG* operon that encodes the minor pilin subunits (Figure 1 and Figure S1). Wei *et al.* examined differences in gene evolution in vaginal bacterial genomes, finding that *Lactobacillus* adhesion-related genes have higher recombination rates compared to their respective genomes and may undergo positive selection (38). Additionally, *Neisseria gonorrhoeae* can undergo recombination of its type IV pilus to avoid antigenic detection (39). Although the *comG*-encoded pilus is not considered a surface adhesin, it is a type IV-like pilus (20), and it is possible that its surface exposure causes selective pressures for change to avoid either immune detection or phage predation.

It remains to be determined the degree to which *L. iners*’ competence is active *in vivo*, however we believe that competence is widespread in the species. The *L. iners* genome is highly mosaic and our analysis (unpublished) of the distribution of genomic islands across the species finds that the presence of one island cannot predict whether that strain will also have a different island (*i.e.,* genomic islands spread horizontally and frequently between *L. iners* isolates independently of one another and with only a weak correlation with respect to vertical inheritance). A 2016 study examined multiple *L. iners’* strains to uncover evolutionary patterns in its core and accessory genomes; it was found that *L. iners* may have over 10 core genes that could have been horizontally acquired from bacterial vaginosis associated bacteria (40). In addition, the study posits that inerolysin could have been horizontally transferred from *Gardnerella* and *Streptococcus* sp. as they both have cytolysins (vaginolysin and streptolysin, respectively) (40). Inerolysin is most closely related to vaginolysin in the cytolysin family (40). It is possible that this cytolysin was transferred via natural competence, but this remains to be answered. It is also unclear if natural competence may enable some *L. iners* strains to predispose an individual towards BV pathogenesis (41).

A previous attempt to genetically modify *L. iners* using natural competence was unsuccessful (9). There, gDNA from a streptomycin resistant mutant was given to the same wildtype strain, which should be identical to the recipient in both sequence and methylation signatures. Differences in strains or cultivation may therefore account for our results. Rampersaud used media consisting of proteose peptone, beef/yeast extract, NaCl, MgSO₄, MnSO₄, K₂HPO₄, glucose and fetal bovine serum (FBS), whereas the SLIM media we used is a modification of Iscove’s Modified Dulbecco’s Medium as the base which primarily comprises free amino acids and simple sugars (21). It is possible that compounds in beef/yeast extract or FBS may be inhibitory towards *L. iners*’ competence, and perhaps that the unsuccessful media was too rich. For example, *S. thermophilus* LMD-9 is only competent in chemically defined media that lacks peptides but has free amino acids instead (42). More specifically, it was found that nutrient rich media, M17, has tryptone that inhibits competence from occurring due to repression of *comX* expression (43). In *Lactococcus lactis*, competence may be regulated by diauxic shifts to secondary sugar sources via derepression of the CcpA (a carbon catabolite regulator) from *comX* (44). Higher pH may also enable derepression of the CovRS stress regulatory system from *comX* (44). It remains to be determined which nutrients or environmental signals enable *L. iners’* competence. A chemically defined SLIM (SLIM-CD) could be used to identify metabolites that impact competence (21).

How *L. iners*’ competence is regulated remains unclear. In *B. subtilis* and *S. pneumoniae* competence is regulated by a quorum-sensing two-component regulatory system (45, 46). However, *S. thermophilus* has a similar regulatory network that may be unlinked to quorum sensing, as it can be induced from early growth to the onset of stationary phase without need for a threshold cell density to occur (43). For *L. lactis*, it is unclear if quorum plays a role, as upstream effectors of two-component CovRS regulating *comX* were not studied (44). *L. iners* may have a temporal window for competence; in a separate test, gDNA added to a 24-hour NVM196S05 subculture did not result in transformants.

Unlike many of the competent Gram-positive bacteria, most *L. iners* strains encode only one single-stranded DNA binding (Ssb) protein, SsbA. *B. subtilis*, *S. thermophilus*, and *S. pneumoniae* encode two *ssb* paralogs, *ssbA* and *ssbB* (47–50). SsbA appears to play a housekeeping role based on its activity in DNA replication in *B. subtilis* (51); it is also an essential gene in *B. subtilis* (52) and *S. pneumoniae* (53). SsbB, in contrast, is thought to be more specialized for competence-related homologous recombination of ssDNA (16). *B. subtilis* and *S. pneumoniae ssbB* mutants display reduced transformation efficiency (up to an order of magnitude), but not total abolishment (52, 54, 55) suggesting some redundancy with SsbA (48, 56, 57). We believe the *L. iners* paralog is indeed ‘SsbA’ as it is encoded proximal to *dnaA* and other chromosomal maintenance genes. NVM025S01 was the only strain in our panel to have an additional Ssb protein (Table S1), but we have yet to test if it enhances transformation.

Further work is necessary to determine why transformation efficiency is not uniform across all *L. iners* strains. It is possible that RM systems play a role in inter-strain transformation (58–61), although we do not yet have sufficient data using RM mutants or varying DNA substrates to delineate their role. The effect of RMs and methylation across competent bacteria is not universal: it appears not to affect transformation of Gram-positive *B. subtilis* (62) and *S. pneumoniae* (63), but does affect transformation in many Gram-negative species including *Acinetobacter baumannii* (58), *Pseudomonas stutzeri* (61), *Helicobacter pylori* (64), and *Neisseria meningitidis* (59). In addition, as incoming DNA is single stranded (and RM systems typically act on dsDNA), it is unclear at which stage of the transformation process they would be effective without generating toxic breaks in the genome (65).

The results presented here will provide an invaluable toolbox for better understanding *L. iners*’ biology. Genetic manipulation of *L. iners*’ using its natural competence will help to uncover what genes and genomic networks may impact transitions to or prevention of BV dysbiosis as well as understanding which genes may be essential for host adherence and interactions.

## ACKNOWLEDGEMENTS

We would like to express gratitude and sincerest thanks to the Kenyan Sex Workers Outreach Program, Dr. Rupert Kaul’s lab and Dr. Rachel Liu for the cervicovaginal samples from which our *L. iners* strains were isolated. This work was funded in part by an IMPACT grant from the Emerging Pandemics and Infections Consortium (EPIC), the Natural Sciences and Engineering Research Council of Canada (NSERC), and the Canadian Institutes for Health Research (CIHR). KYC, DS, and JRC were funded by Ontario Graduate Scholarships. KYC was also funded by a Canada Graduate Scholarship (CGS-M). We would also like to thank Dr. Landon Getz for his insightful discussions on the bioinformatics and genomic data help as well as Dr. Dixon Ng and Trevor Bell for allowing us to use their computational resources for the base-calling.

## Supplemental Material

**Figure S1.**
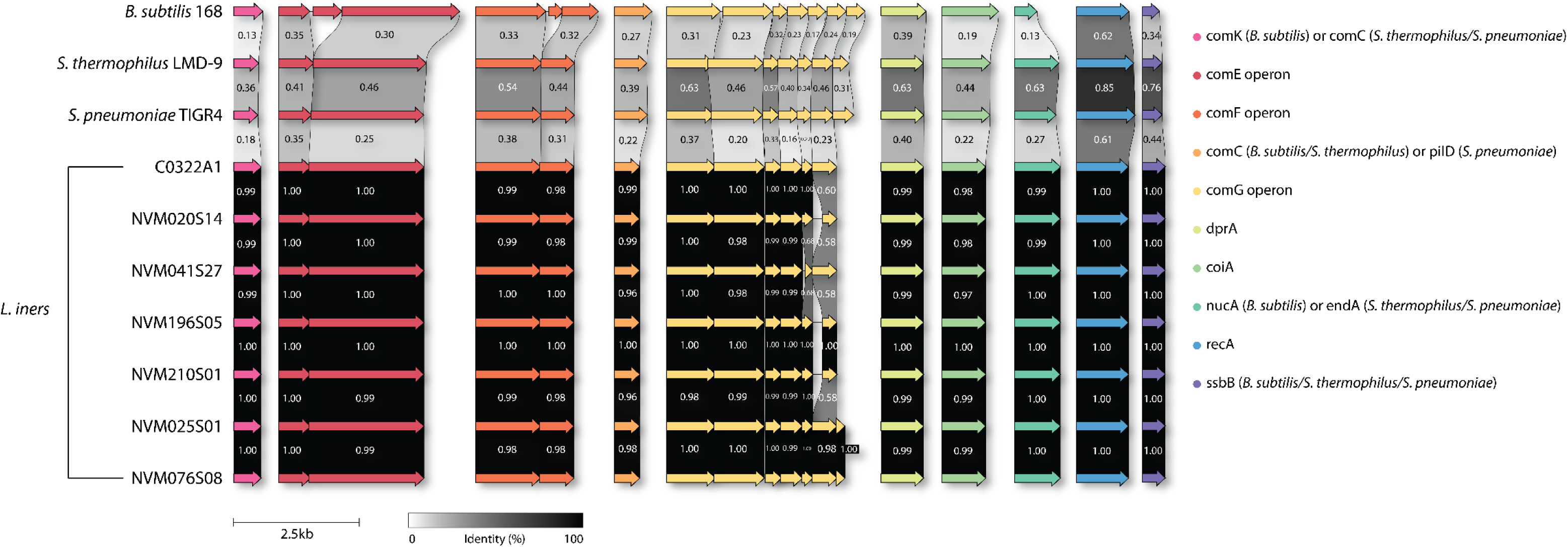
Genomic analysis and detailed identity scores of putative *L. iners* competence operons to known competent model bacteria. *L. iners* strains from Kenyan sex workers were examined for competence genes that may be homologous to model Gram-positive naturally competent bacteria (*B. subtilis*, *S. thermophilus*, *S. pneumoniae*).

**Figure S2.**
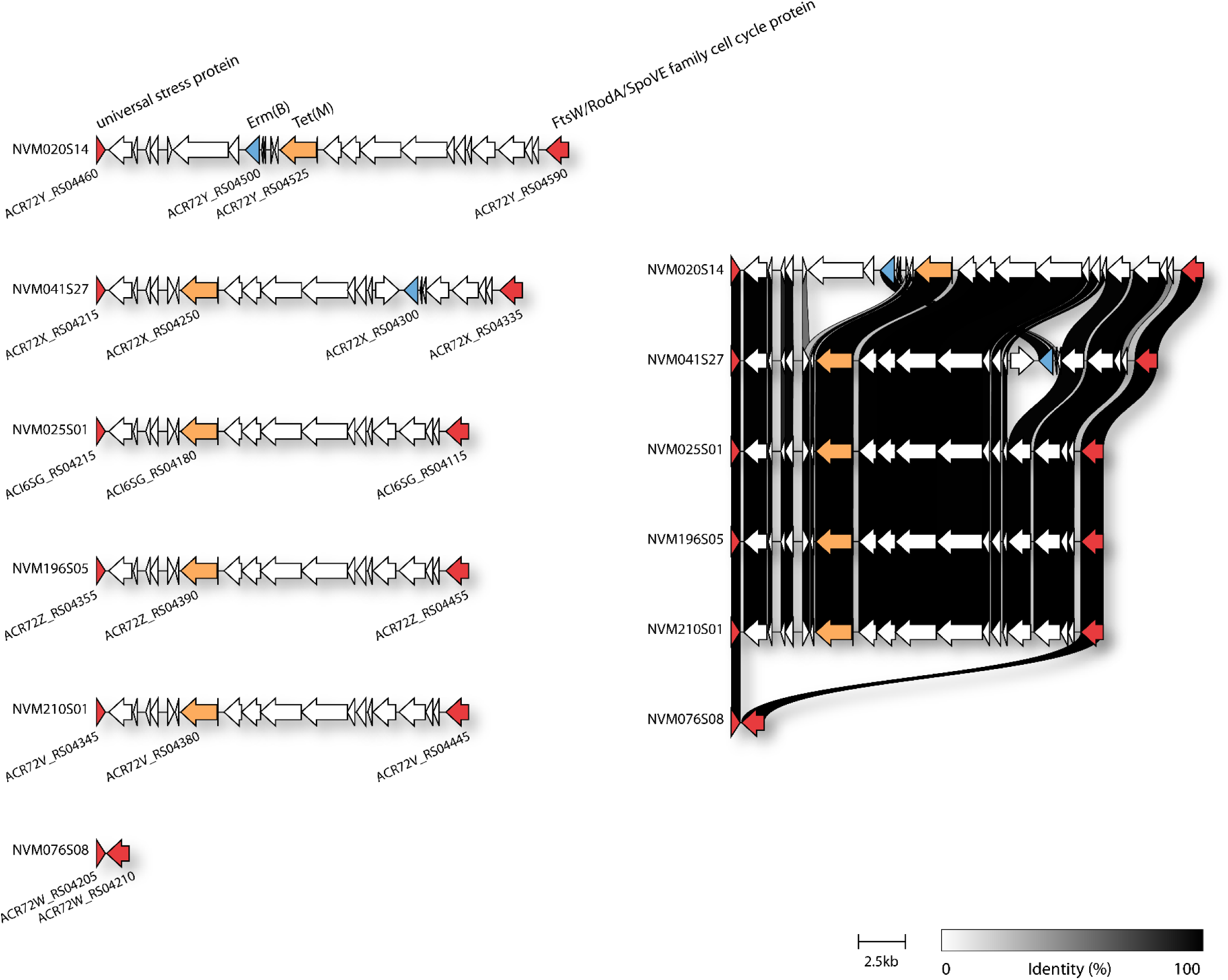
RefSeq locus tags and homology for Tn916 transposable island across *L. iners* strains.

**Figure S3.**
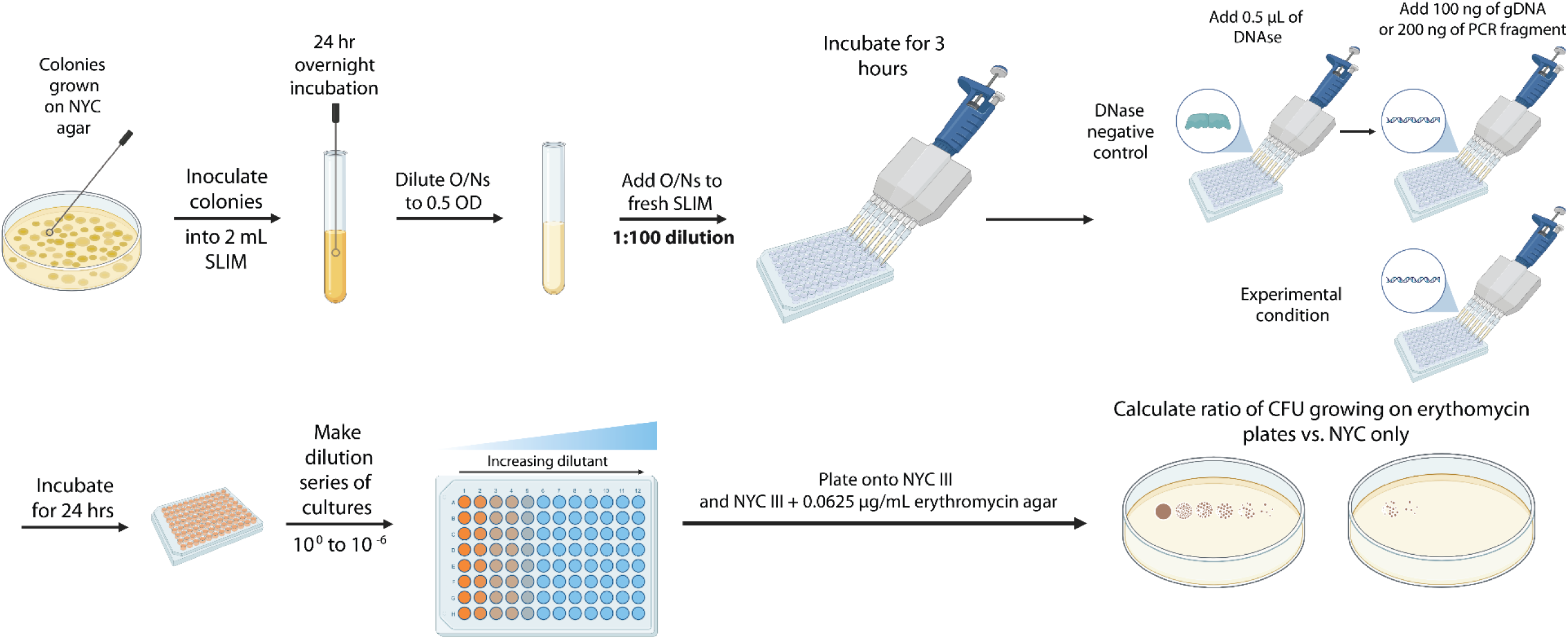
Overview of transformation protocol for *L. iners*.

**Figure S4.**
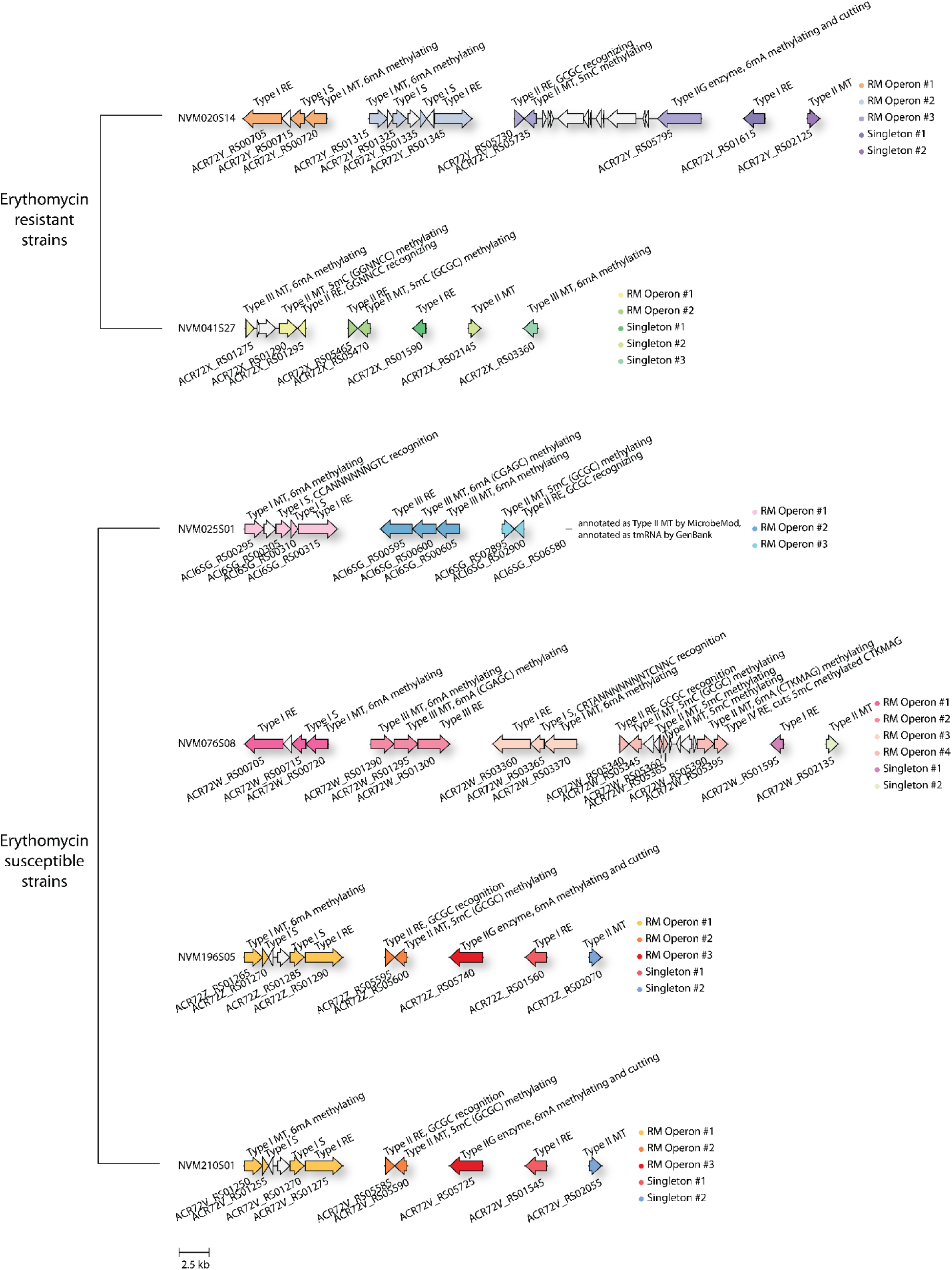
Restriction-modification systems for *L. iners* strains. RM systems in *L. iners* genomes were searched for using MicrobeMod; RM operons generally indicate restriction enzyme (RE) with a cognate methylase (MT) and/or specificity (S) enzyme; singletons indicate incomplete RM system.

**Figure S5.**
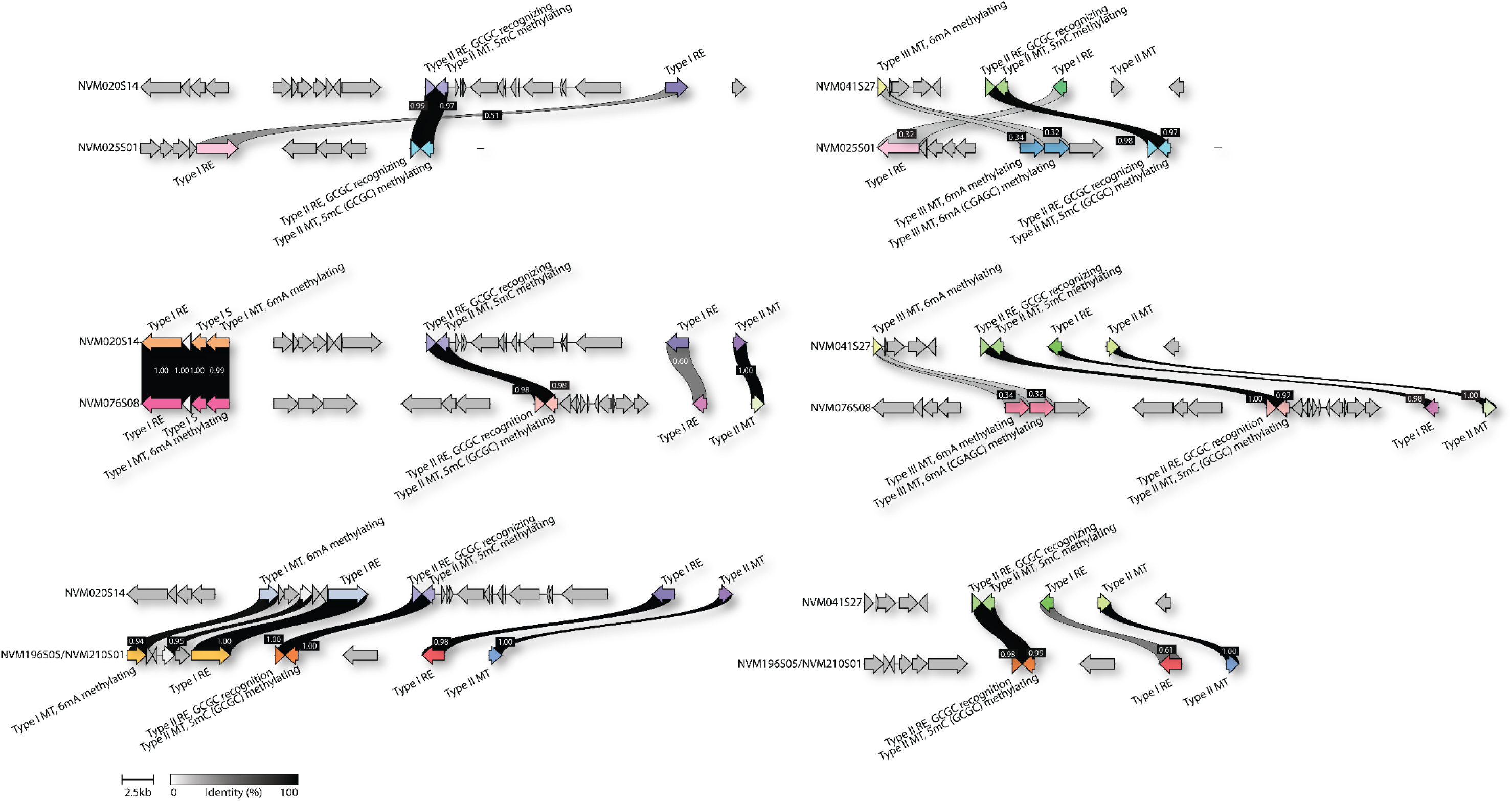
Homology of restriction systems between *L. iners* strains. Percent shared nucleotide identity between enzymes shown in black linker boxes.

**Figure S6.**
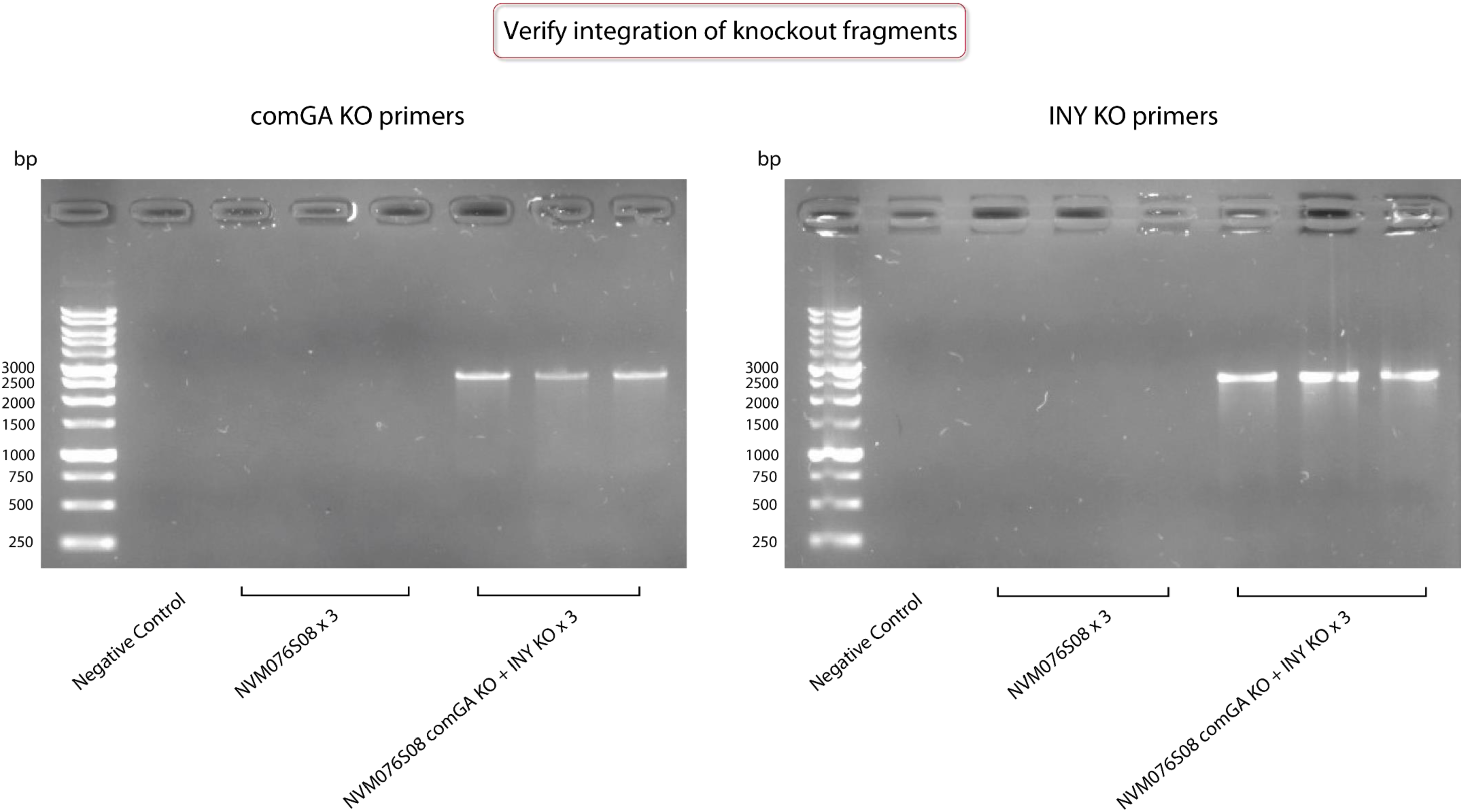
Confirmation of generation of an NVM076S08 *comGA^−^ iny^−^* double knockout. Left gel with *comGA* integration primers (expected band size 2663 bp), right gel with *INY* KO integration primers (expected band size 2534 bp). Negative control has no colonies added.

**Table S1.** Genome coordinates and RefSeq annotations of competence operons in Gram-positive models and *L. iners* strains. Coordinates are genomic coordinates for entire operon (e.g., entire comG operon). Locus tag is RefSeq ID for each gene. Gene symbol and product is from RefSeq annotation. Protein ID is GenBank accession code for protein. Feature Start and Feature End indicate genomic coordinates for an individual gene. Feature Type is coding sequence (CDS). Strand indicates +ve or -ve sense (i.e., how operons are organized relative to one another in terms of which DNA strand encodes for operon). **See attached Excel File**

**Table S2.**
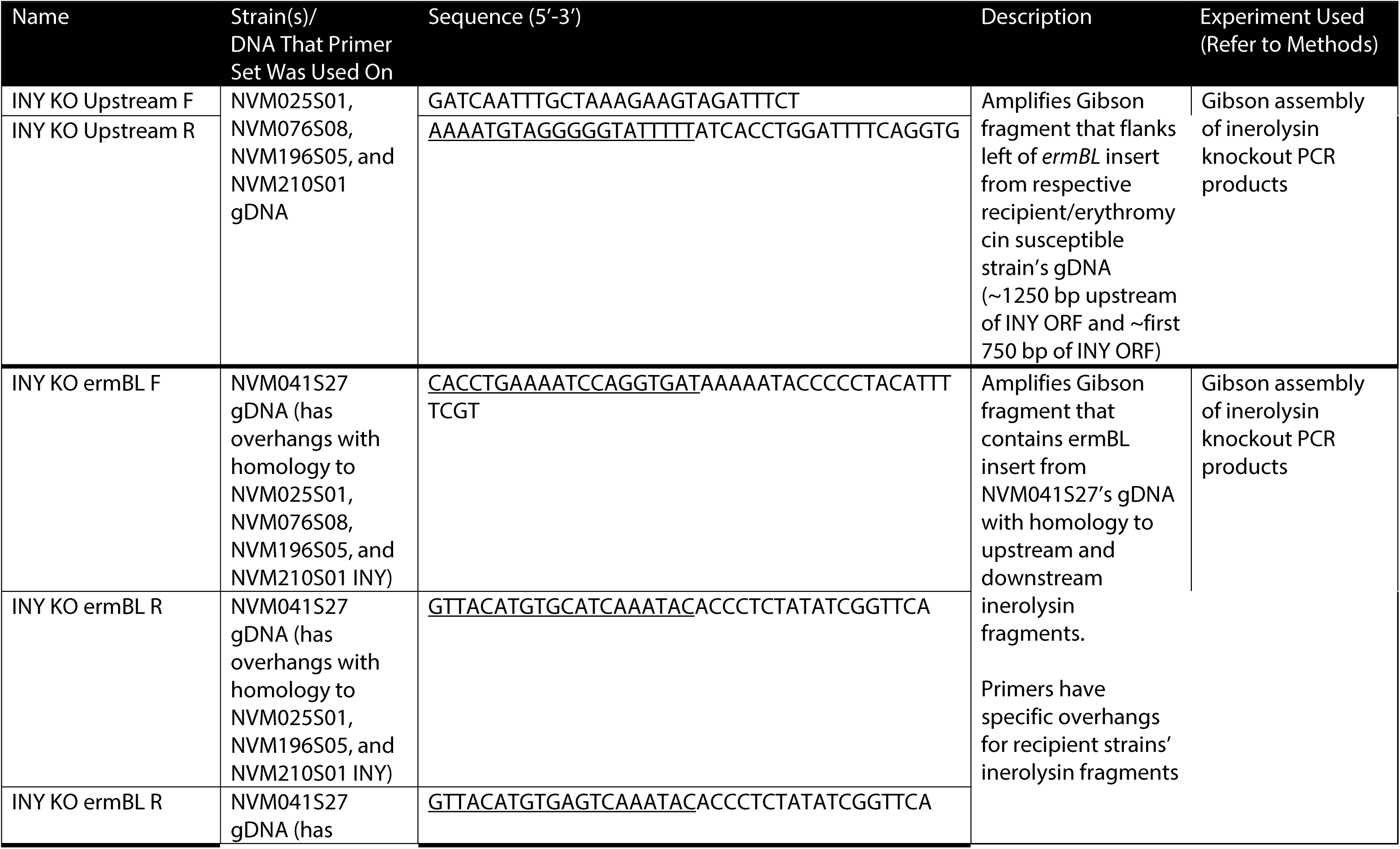
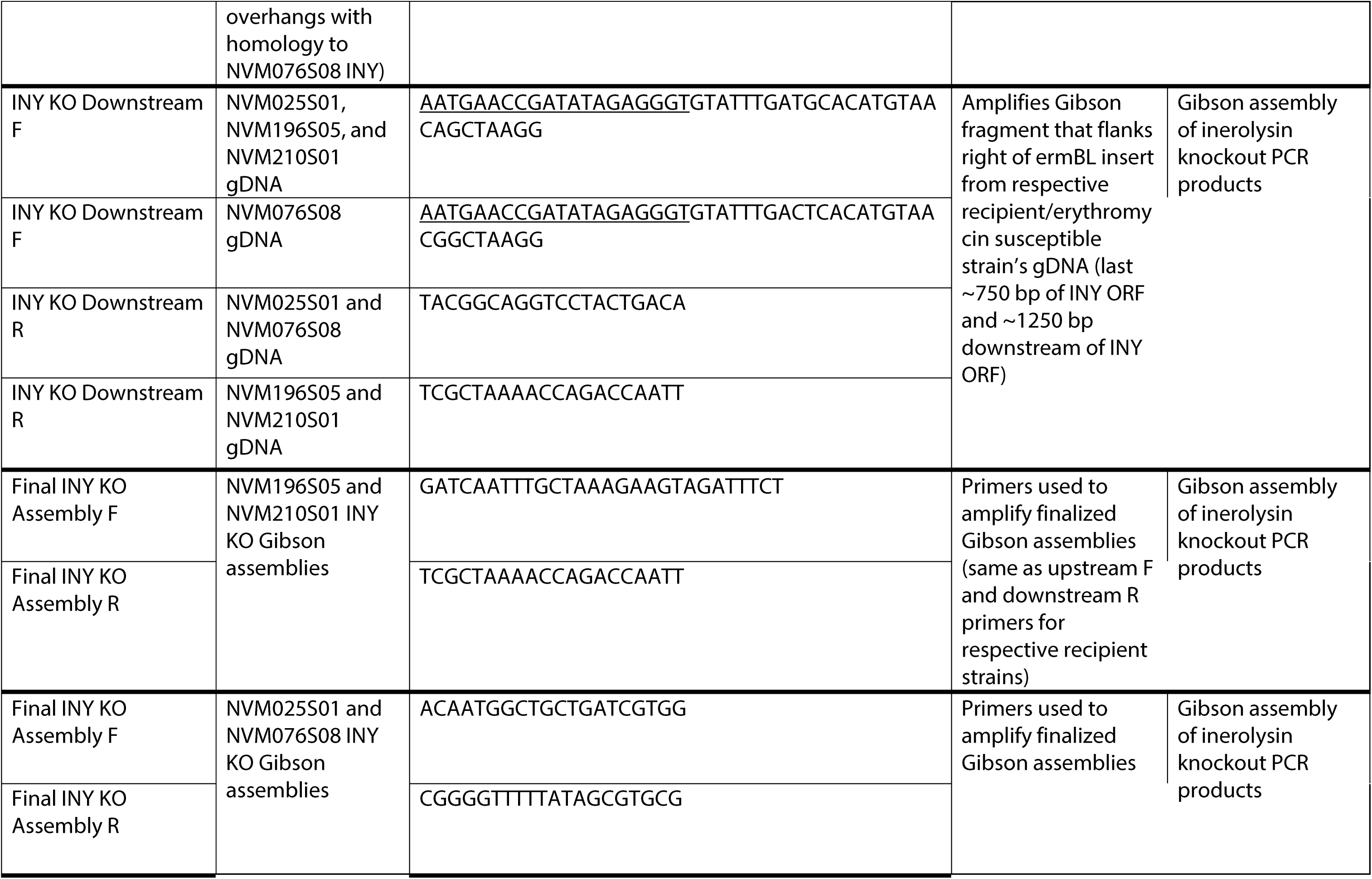
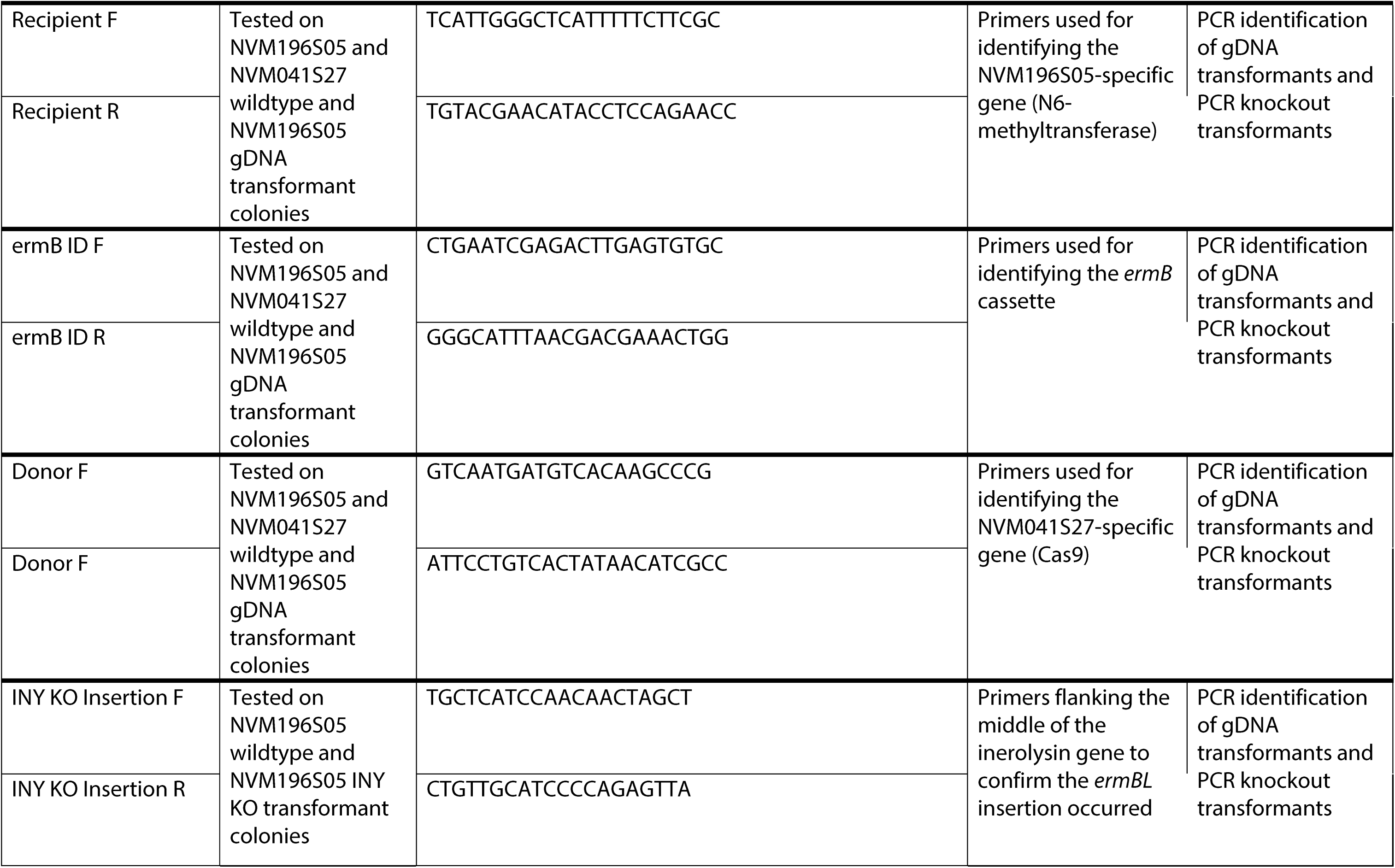
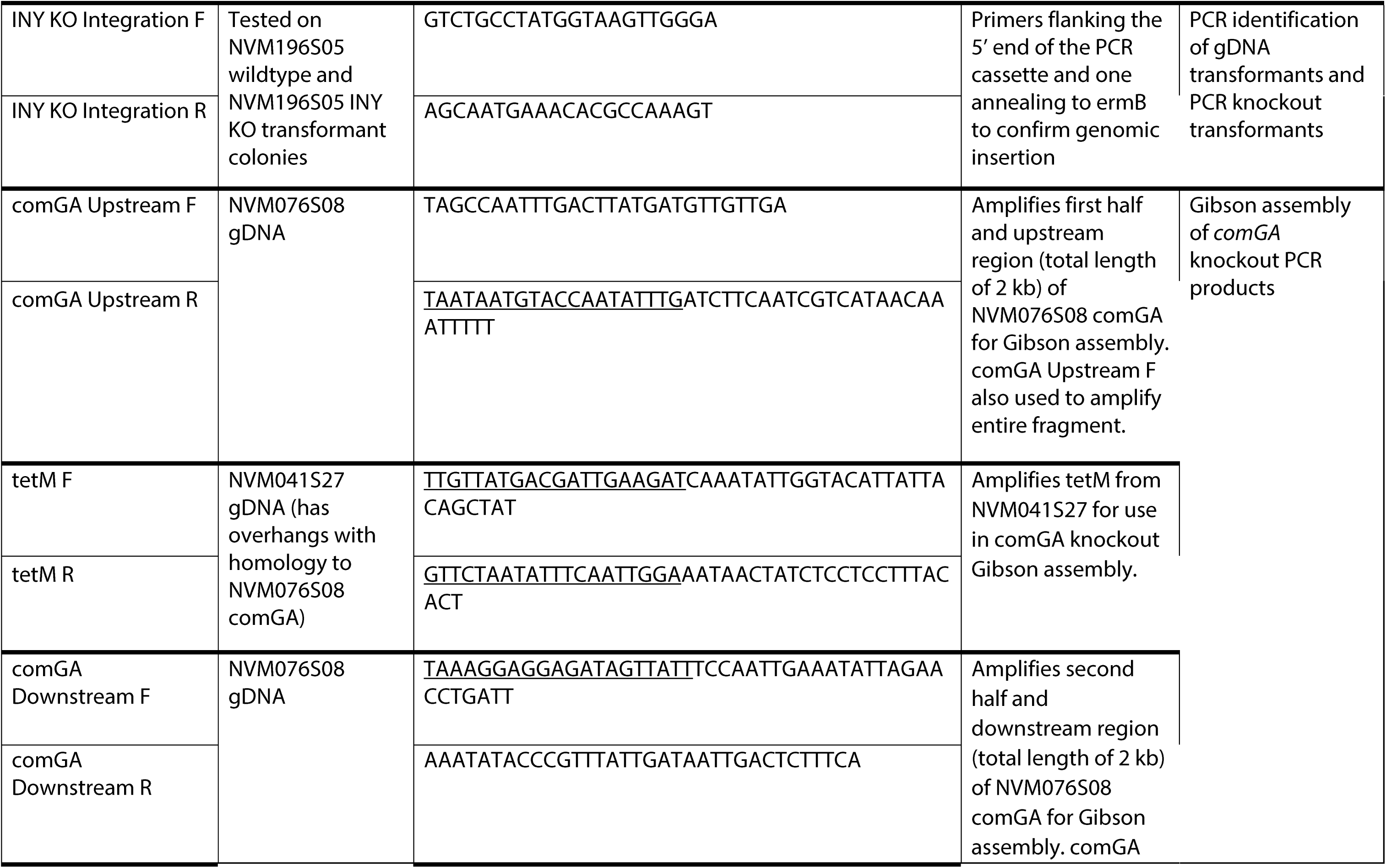
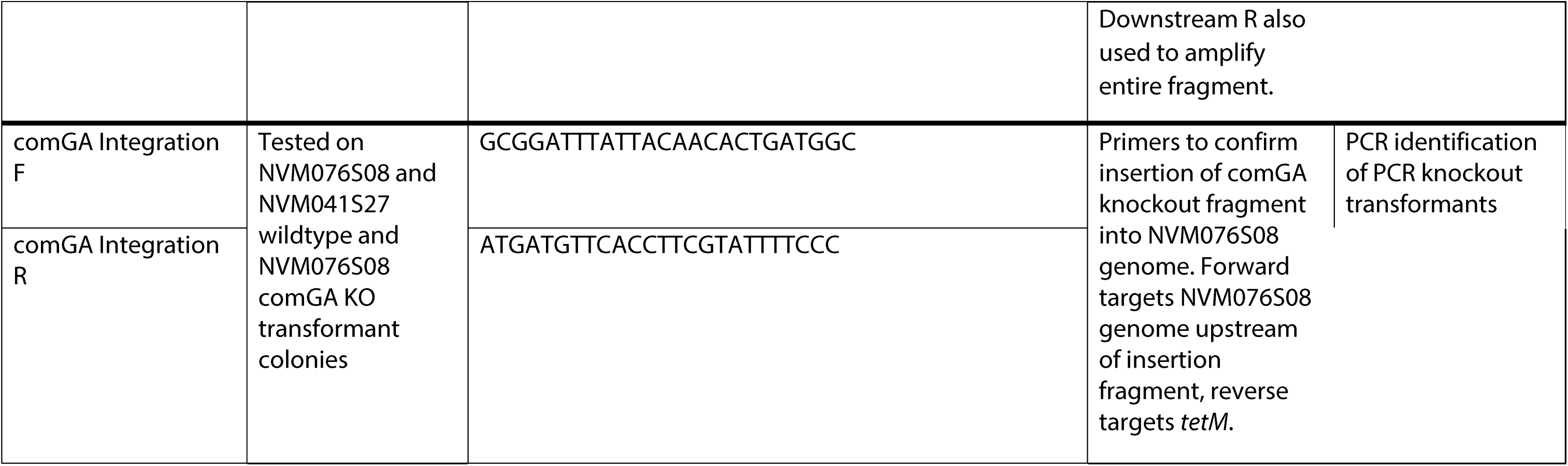
Primers used. Overhangs used for Gibson homology regions underlined.

**Table S3.**
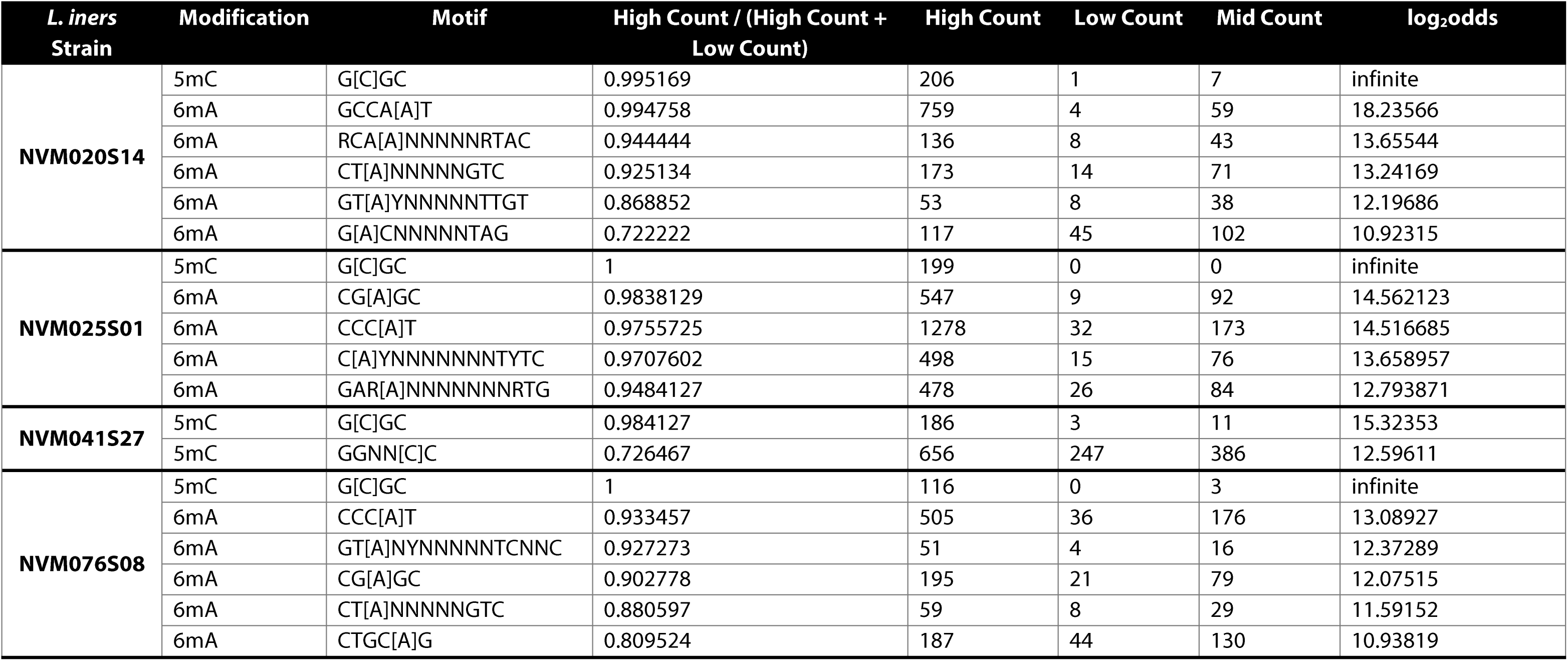
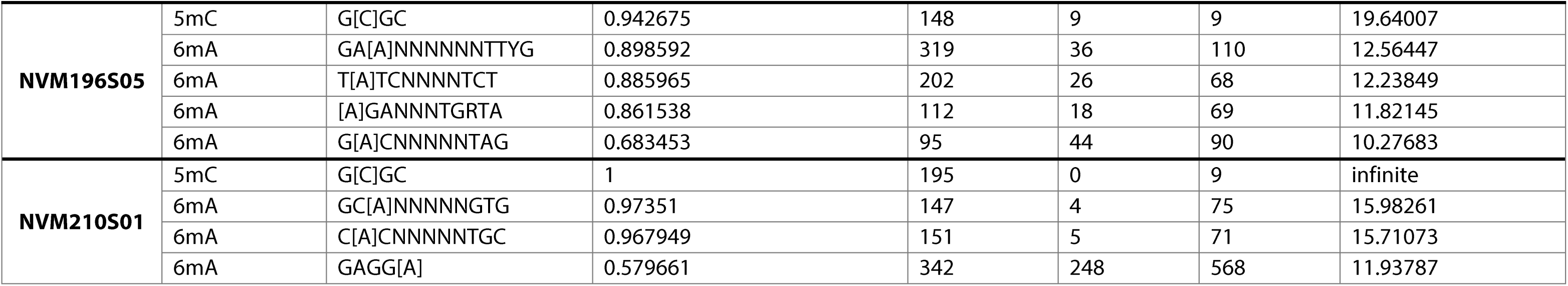
Predicted methylation motifs in Nanopore-sequenced *L. iners* strains. Methylation sites were identified from Nanopore sequencing data using Modkit. The Modification column indicates the type of base modification: 5mC (5-methylcytosine) or 6mA (N6-methyladenine). In the Motif column, brackets [] indicate the methylated position. For each motif, Modkit calculates a per-site fraction modified, defined as N_mod / (N_canonical + N_mod + N_other_mod), where N_mod is the number of base calls passing modification filters, N_canonical is the number not passing modification filters, and N_other_mod is the number passing filters for a different modification of the same base. Each genomic site matching a given motif is then classified based on this fraction: High (>= 0.6), Low (< 0.2), or Medium (between 0.2 and 0.6). The High Count, Low Count, and Mid Count columns report the number of genomic sites for that motif falling into each category. The High Count / (High Count + Low Count) column gives the proportion of classifiable sites that are in the High category. The log2odds column is the log2 of the odds of a site being classified as High rather than not High.

